# *Symbiodinium* functional diversity and clade specificity under global change stressors

**DOI:** 10.1101/190413

**Authors:** S.W. Davies, J.B. Ries, A Marchetti, Rafaela Granzotti, K.D. Castillo

## Abstract

Coral bleaching episodes are increasing in frequency, demanding examination of the physiological and molecular responses of corals and their *Symbiodinium* to climate change. Here we quantify bleaching and *Symbiodinium* photosynthetic performance of *Siderastrea siderea* from two reef zones after long-term exposure to thermal and CO_2_-acidification stress. Molecular response of *in hospite Symbiodinium* to these stressors was interrogated with RNAseq. Elevated temperatures reduced photosynthetic efficiency, which was highly correlated with bleaching status. However, photosynthetic efficiencies of forereef symbionts were more negatively affected by thermal stress than nearshore symbionts, indicating greater thermal tolerance in nearshore corals. At control temperatures, CO_2_-acidification had little effect on symbiont physiology, although forereef symbionts exhibited greater photosynthetic efficiencies than nearshore symbionts. Transcriptome profiling revealed that *S. siderea* were dominated by clade C *Symbiodinium*, except under thermal stress, which caused shifts to thermotolerant clade D. Comparative transcriptomics of conserved genes across symbiotic partners revealed few differentially expressed *Symbiodinium* genes when compared to corals. Instead of responding to stress, clade C transcriptomes varied by reef zone, with forereef *Symbiodinium* exhibiting enrichment of genes associated with photosynthesis. Our findings suggest that functional variation in photosynthetic architecture exists between forereef and nearshore *Symbiodinium* populations.

## INTRODUCTION

Dinoflagellates are ubiquitous unicellular algae that can occur as free-living individuals, endosymbionts, or parasites across a wide variety of marine organisms. The most recognized among endosymbionts is the genus *Symbiodinium*, which establish obligate relationships with many reef-building corals (Trench and Blank 1987). Photosynthetically-derived nutrients translocated from *Symbiodinium* to the host can comprise up to 100% of the coral energy budget (Muscatine and Cernichiari 1969, Muscatine 1990), allowing for prolific coral growth in shallow oligotrophic tropical waters. In turn, *Symbiodinium* gain a protected microenvironment enriched in light, inorganic nutrients, and dissolved inorganic carbon as provided by the host (Barott et al 2015, Muscatine and Cernichiari 1969, Yellowlees et al 2008). Although initially considered a single species, advancements in genetics have uncovered nine divergent clades within *Symbiodinium*, designated A-I (Coffroth and Santos 2005, Pochon and Gates 2010, Thornhill et al 2014), with growing evidence for additional within-clade genetic diversity (Davies et al 2016b, Howells et al 2012, Pettay and Lajeunesse 2013, Thornhill et al 2017).

Increasing atmospheric *p*CO_2_ has reduced seawater pH, which impairs reef accretion (Hoegh-Guldberg et al 2007), and has increased sea surface temperatures, which can cause a breakdown of the coral-algal symbiosis—termed coral bleaching (Baker 2003, Hoegh-Guldberg 1999, Hoegh-Guldberg and Bruno 2010, Weis 2008). During bleaching, the coral is deprived of symbiont-derived organic carbon, which may lead to reduced growth or mortality. Bleaching events have been recognized as the primary driver of recent global reef declines and have been increasing in frequency and severity (Hughes et al 2017, Pandolfi et al 2011). However, the rich genetic diversity of *Symbiodinium* often reflects their functional diversity, which could confer resilience to changing environments. For example, within-host *Symbiodinium* communities can shift after bleaching disturbances (Baker 2001, Jones et al 2008, Kemp et al 2014, Silverstein et al 2015, Thornhill et al 2006), which can strongly influence coral gene expression and future bleaching susceptibility (DeSalvo et al 2010a, Howells et al 2012, Jones and Berkelmans 2010). In order to predict coral reef responses to global change, there is an urgent need for a more comprehensive understanding of how coral-*Symbiodinium* associations respond to ocean warming and acidification.

Studies of *Symbiodinium* physiology document a wide array of stress responses to environmental change, including impairment/inactivation of photosynthesis at high temperatures (Iglesias-Prieto et al 1992, Iglesias-Prieto and Trench 1994), increased production of reactive oxygen species (ROS) and antioxidant activity in response to thermal and UV exposure (Gardner et al 2017, Lesser 1996, Suggett et al 2008), and reduced symbiont pigment concentrations at low pH (Tremblay et al 2013). In contrast to these strong physiological responses, there is mounting evidence that *Symbiodinium* lack strong transcriptional responses to global change stressors, which starkly contrasts their host’s transcriptional response (Barshis et al 2014, Leggat et al 2011). Leggat et al. (2011) found that *Symbiodinium* exhibit few changes in expression of stress response genes under thermal stress when compared to these same genes in their coral host. A lack of transcriptomic responses has also been observed in both *Symbiodinium* D2 and C3K following heat exposure (Barshis et al 2014), which contrasts wide-spread transcriptomic shifts observed in the coral host in response to heat stress (Barshis et al 2013). Despite the unresponsiveness of *Symbiodinium* gene expression, transcriptome profiles were clade-specific, regardless of experimental treatment, supporting prior evidence of functional differences amongst clades (Barshis et al 2014). Within clade functional variation has also been documented in clade B strains: when transcriptomes were compared, lineage-specific differences in expression were observed (Parkinson et al., 2016), providing additional evidence that substantial functional variation exists amongst *Symbiodinium* clades.

Clade C strains are the most derived *Symbiodinium* lineage and exhibit higher within-clade diversity when compared to other, more basal clades (Lesser et al 2013, Pochon et al 2006, Pochon and Gates 2010). Understanding how this diversity affects a coral’s response to global change stressors is of great interest because clade-specific responses to stress are well documented. Transcriptomes of two divergent clade C1 lineages previously shown to exhibit distinct thermal tolerances (Howells et al 2012) revealed divergent expression patterns in response to heat stress, providing insights into how functional variation could lead to variation in holobiont (coral + *Symbiodinium*) thermal tolerance (Levin et al 2016). Given that coral *host-Symbiodinium* interactions ultimately determine how the holobiont will respond to global change, it is critical to determine how each symbiotic partner responds to multiple stressors. However, few studies have explored whole transcriptome responses of different *Symbiodinium* populations to ocean warming and acidification *in hospite* and even fewer have compared the responses of both symbiotic partners in parallel.

Here, we exposed the resilient and ubiquitous Caribbean reef-building coral *Siderastrea siderea* and its clade C *Symbiodinium* from two different reef zones (nearshore and forereef) to thermal (*T* = 25, 28, 32°C) and CO_2_-induced acidification (*p*CO_2_ = 324, 477, 604, 2553 μatm) stress for 95 days. Our previous work has shown that both elevated temperature and *p*CO_2_ elicited strong but divergent responses of the host’s transcriptome (Davies et al 2016a) with increased temperatures substantially reducing calcification rates, with smaller declines observed in response to elevated *p*CO_2_ (Castillo et al 2014). Here, we build on these studies by quantifying holobiont bleaching response, *Symbiodinium* photosynthetic efficiencies (F_v_/F_m_), and transcriptomic responses of *Symbiodinium in hospite*. We disentangle the response of the holobiont by isolating the responses of the host coral and *Symbiodinium* to thermal and acidification stress, which advances our understanding of their potential ecological trajectories in response to these global change stressors.

## MATERIALS AND METHODS

### Experimental design

Physiological measurements, transcriptomes and gene expression analyses presented here build upon experiments previously published in Castillo et al. (2014) and Davies et al. (2016). Briefly, whole *S. siderea* colonies from nearshore (N=6) and forereef (N=6) reef zones were collected from the Belize Mesoamerican Barrier Reef (total colonies = 12) in July 2011 and transported to the University of North Carolina at Chapel Hill (UNC) Aquarium Research Center. At UNC, whole coral colonies were sectioned into 18 fragments, which were maintained across six experimental treatments (N=3 tanks per treatment) that were illuminated with 250 μmol photons m^-2^ s^-1^ on a 12-hour light-dark cycle for 95 days. Partial pressures of CO_2_ corresponded to near-pre-industrial (324 ± 89 μatm, 28.14 ± 0.27 °C), near-present-day (*p*CO_2_ 477 ± 83 μatm, 28.16 ± 0.24 °C), predicted end-of-century (604 ± 107 μatm, 28.04 ± 0.28 °C), and an extreme mid-millennium scenario (2553 ± 506 μatm, 27.93 ± 0.19°C). Seawater temperatures approximated monthly minimum (25.01 ± 0.14 °C, *p*CO_2_ 515 ± 92 μatm), mean (28.16 ± 0.24 °C, *p*CO_2_ 477 ± 83 μatm), and maximum seawater temperatures (32.01 ± 0.17 °C, *p*CO_2_ 472 ± 86 μatm) over the interval from 2002 to 2014 (Castillo and Helmuth 2005, Castillo and Lima 2010, Castillo et al 2012). Upon completion of the experiment, holobiont tissue (coral + symbiont) was extracted by water pick, immediately preserved in RNAlater, and stored at −80°C until RNA was extracted.

### Symbiodinium *physiology: tissue bleaching and photosynthetic efficiency*

Coral fragments were photographed at the end of the 95-day experiment prior to tissue extraction. To quantify coral color differences associated with reduced chlorophyll and symbiont densities, Adobe Photoshop balance exposures were standardized across photographs using a white standard. Total red, green, blue and sums of all color channel intensities (red, green, blue) were calculated for 10 quadrats of 25 x 25 pixels within each coral fragment as a measure of brightness, with higher brightness indicating a reduction in algal pigment (i.e., reduced symbiont density or bleaching). This analysis was performed following Winters et al (2009) using the MATLAB macro “AnalyzeIntensity” and resulting data were inverted so that higher values represented increased coral pigmentation.

Photosynthetic efficiencies of the symbionts’ photosystem II (F_v_/F_m_) were quantified for each experimental fragment using pulse amplitude modulated fluorometry (PAM; Diving-PAM, Waltz). Reductions in F_v_/F_m_ values are indicative of sustained damage to the photosystem. F_v_/F_m_ measurements were collected just prior to completion of the experiment and were made following dark adaptation one hour after sunset, which was when the lights turned off (n=3 measurements per coral fragment).

All statistical analyses were implemented in R (R Development Core Team 2015) using the ANOVA function based on log-transformed sum of all color channels and F_v_/F_m_ data. The sum of all channels was chosen as the best color metric proxy since these data correlated best with F_v_/F_m_ (Adjusted *r*^*2*^: 0.7366, *p*-value<0.001). Experimental treatment and reef zone were modeled as fixed effects and significant differences across levels within factors were evaluated using post-hoc Tukey’s HSD tests. All assumptions of parametric testing were explored using diagnostic plots in R.

### *Transcriptome assembly, annotation and* Symbiodinium *identification*

Detailed descriptions of sequence library preparations, holobiont transcriptome assembly and separation of host and symbiont contigs can be found in Davies *et al*. (2016a). Briefly, RNA was isolated from 94 holobiont fragments and pooled by reef zone within each experimental treatment, yielding a total of twelve sequencing libraries (Table 1), each of which contained RNA from at least six holobionts. Since each coral holobiont can host a community of *Symbiodinium*, RNA pooling is not expected to yield any additional complications compared to preparing each holobiont independently. Pooled libraries were prepared and sequencing was performed using four lanes of *Illumina HiSeq* 2000 at the University of North Carolina High Throughput Sequencing Facility, which yielded paired-end (PE) 100bp reads (Table 1).

**Table 1.** Summary of RNA libraries, including the number of holobionts from which RNA was pooled (’Pool’), reef zone where holobionts were collected (forereef = ‘FR’ and nearshore = ‘NS’), temperature (°C) ±SD, *p*CO_2_ (μatm) ±SD, raw 100bp paired-end reads (‘PE Reads’), mapped holobiont reads (‘Mapped’) and total number of Symbiodinium-specific counts (‘Symbiont’). Shaded samples were excluded from analyses due to low counts as a result of bleaching for a total of 235.1 million reads used in downstream analyses.

| Sample | Pool | Reef zone | Temp (°C) | $p\text{CO}_2$ ($\mu\text{atm}$) | PE Reads ($10^6$) | Mapped ($10^6$) | Symbiont ($10^6$) |
| --- | --- | --- | --- | --- | --- | --- | --- |
| FR_P324 | 6 | FR | 28.14 ± 0.27 | 324 ± 89 | 63.6 | 59.2 | 24.1 |
| FR_P604 | 6 | FR | 28.04 ± 0.28 | 604 ± 107 | 77.1 | 72.3 | 16.8 |
| FR_P2553 | 7 | FR | 27.93 ± 0.19 | 2553 ± 506 | 48.1 | 44.7 | 17.1 |
| NS_P324 | 6 | NS | 28.14 ± 0.27 | 324 ± 89 | 88.3 | 82.4 | 35.0 |
| NS_P604 | 8 | NS | 28.04 ± 0.28 | 604 ± 107 | 48.5 | 45.3 | 10.5 |
| NS_P2553 | 6 | NS | 27.93 ± 0.19 | 2553 ± 506 | 69.2 | 64.6 | 23.3 |
| FR_T25 | 11 | FR | 25.01 ± 0.17 | 515 ± 92 | 50.9 | 47.4 | 20.5 |
| FR_Control | 14 | FR | 28.16 ± 0.24 | 477 ± 83 | 63.2 | 58.8 | 24.3 |
| FR_T32 | 9 | FR | 32.01 ± 0.17 | 472 ± 86 | 50.0 | 47.2 | 1.2 |
| NS_T25 | 7 | NS | 25.01 ± 0.17 | 515 ± 92 | 100.0 | 93.1 | 39.9 |
| NS_Control | 7 | NS | 28.16 ± 0.24 | 477 ± 83 | 56.7 | 52.7 | 23.6 |
| NS_T32 | 7 | NS | 32.01 ± 0.17 | 472 ± 86 | 54.7 | 51.5 | 3.9 |
| **TOTAL** |  |  |  |  | **770.3** | **719.2** | **235.1** |

Transcriptome *de novo* assembly of over 770 million PE reads was completed using Trinity (Grabherr et al 2011) and resulting contigs were assigned as either coral-or *Symbiodinium-derived* contigs using a translated blast search (tblastx) against host and symbiont specific databases (Davies et al 2016a). Symbiodinium-specific contigs were annotated by BLAST sequence homology searches against UniProt and Swiss-Prot NCBI NR protein databases with an e-value cutoff of e^-5^ (Consortium 2015). Annotated sequences were then assigned to Gene Ontology (GO) categories. Transcriptome contiguity analysis (Martin and Wang 2011) and Benchmarking Universal Single-Copy Orthologs v2 (BUSCO; Simao et al 2015) were used to assess *Symbiodinium* transcriptome quality and completeness.

### Symbiodinium *community composition*

To compare *Symbiodinium* expression across reef zones and experimental treatments, it was first necessary to determine which *Symbiodinium* clades were present in each library. Reads were trimmed with Fastx_toolkit (<20 bp length cutoff and bp quality score >20) and resulting quality filtered reads were then mapped to an ITS2-specific database (Franklin et al 2012) using Bowtie2.2.0 with the ‒a flag to search all possible alignments (Langmead and Salzberg 2012). All possible alignments were identified and then the read was quantified in the analysis if all possible alignments for that read mapped to a single clade. This analysis determined that 96.7 to 100% of mapped reads from 10 of the 12 libraries exclusively aligned to a single *Symbiodinium* clade, with remaining libraries (NS_T32 and FR_T32) hosting divergent communities. However, these two libraries hosting divergent communities were excluded from downstream expression analysis due to the temperature-induced bleaching in these treatments yielding low concentrations of algal cells, which ultimately led to insufficient quantities of mapped *Symbiodinium* reads (1.2 and 3.9 million).

### *Mapping and differential expression analysis for clade C* Symbiodinium

Raw reads across samples ranged from 48.1 to 100.0 million PE 100bp sequences. Quality filtered reads were mapped to the holobiont transcriptome (*Siderastrea siderea* + clade C *Symbiodinium*) and *Symbiodinium* mapped reads ranged from 10.5 (NS_P604) to 39.9 (NS_T25) million (Table 1). Differential gene expression analyses were performed on raw counts with *DESeq2* (v. 1.6.3; Love et al 2014) in R (v. 3.1.1; R Development Core Team 2015) using the model: design ~ reef zone + treatment. Counts were normalized for size factor differences and pairwise contrasts were computed for each treatment relative to the control and between the two reef zones. A contig was considered significantly differentially expressed if it had an average basemean expression >5 and an FDR adjusted *p*-value < 0.05 (Benjamini and Hochberg 1995). Raw counts were then *rlog* normalized, a principle coordinate analysis was performed and the *adonis* function tested for overall expression differences across treatments and reef zones (Oksanen et al 2013). Gene expression heatmaps with hierarchical clustering of expression profiles were created with the *pheatmap* package in R (Kolde 2012). Gene ontology (GO) enrichment analysis was performed using the GO_MWU method, which uses adaptive clustering of GO categories and Mann-Whitney U tests (Voolstra et al 2011) based on a ranking of signed log p-values (Dixon et al 2015). Results were plotted as a dendrogram, which traces the level of gene sharing between significant categories and lists the proportion of genes in the dataset with raw p-values < 0.05 relative to the total number of genes within the entire expression dataset.

### Expression comparison across holobiont partners

In order to compare expression across symbiotic partners, a subset of highly conserved genes (HCG) from the coral host (*Siderastrea siderea*) and clade C *Symbiodinium* were mined based on conserved gene annotation (N=2,862 genes). Differential gene expression analyses were performed with *DESeq2* using only data from the HCG set (design ~ reef zone + partner.treatment). Counts were normalized for size factor differences and pairwise contrasts were computed for each treatment relative to the control for each symbiotic partner. A contig was considered significantly differentially expressed if it had an average basemean expression > 5 and an FDR adjusted *p*-value < 0.05 (Benjamini and Hochberg 1995). Significant DEGs were then compared across symbiotic partners. In order to confirm that *Symbiodinium* gene expression results were not confounded by strong reef zone differences in expression, heatmaps were generated on the HCG panel that was significantly differentially expressed in the host but not the symbiont.

## RESULTS

### *Reef zone variation in* Symbiodinium *physiology*

Long-term thermal stress treatments resulted in significantly reduced F_v_/F_m_ of *Symbiodinium* (p<0.001; Fig. 1A), which was highly correlated with bleaching status (p<0.001, r^2^=0.74; Fig. 1B). Interestingly, *Symbiodinium* originating from nearshore habitats had significantly higher F_v_/F_m_ under thermal stress when compared to forereef *Symbiodinium* (Tukey’s HSD p=0.008), suggesting increased resilience to warmer temperatures. However, nearshore *Symbiodinium* only exhibited significantly higher F_v_/F_m_ compared to forereef symbionts in 32°C treatments; in all other experimental treatments, forereef symbionts had constitutively higher photosynthetic efficiencies (Fig. 1A; *p*<0.001).

**Figure 1:**
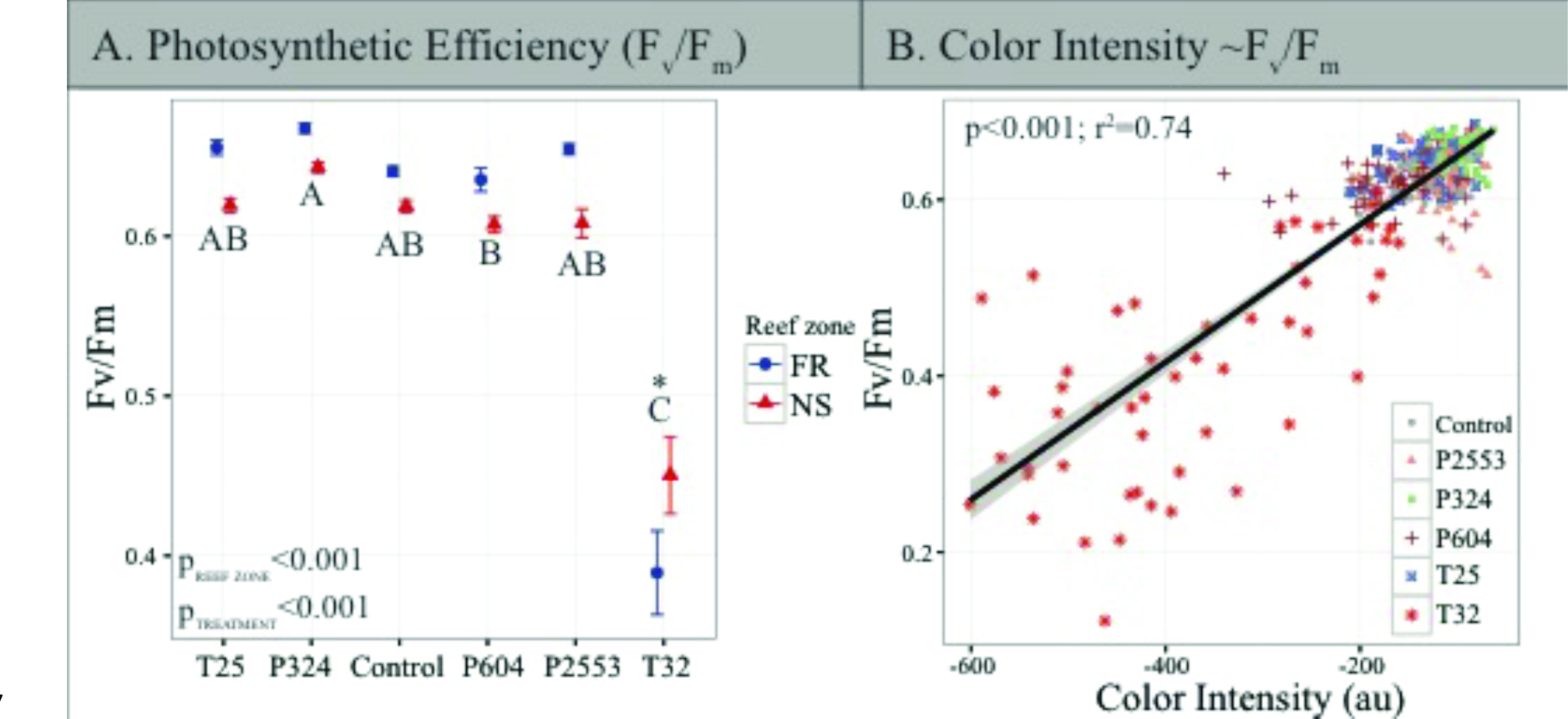
Physiological measurements of *in hospite Symbiodinium*. (A) Photosynthetic efficiency of *Symbiodinium* photosystem II (PSII) (F_v_/F_m_) after 95 days in experimental treatments. Blue circles: forereef population average (±SE); red triangles: nearshore population average (±SE). Asterisk (*) indicates a significant interaction between treatment and reef zone. Treatment pairs lacking a letter in common indicate a statistically significant difference between those treatments pursuant to Tukey’s Honest Significant Difference tests. High temperatures reduce F_v_/F_m_, however, symbonts from nearshore environments are less affected than those from the forereef, while forereef symbionts exhibit higher F_v_/F_m_ in all other treatments. (B) Maximum F_v_/F_m_ of *Symbiodinium* vs. mean coral bleaching status after 95 days in experimental treatments (T25= 25.01 ± 0.14 °C, 515 ± 92 μatm; Control= 28.16 ± 0.24 °C, 477 ± 83 μatm; P324= 28.14 ± 0.27 °C, 324 ± 89 μatm; P604= 28.04 ± 0.28 °C, 604 ± 107 μatm; P2553= 27.93 ± 0.19°C, 2553 ± 506 μatm; T32= 32.01 ± 0.17 °C, 472 ± 86 μatm), as measured by the sum of all color intensities in the red, green and blue channels in standardized coral photographs across all treatments. This relationship demonstrates that F_v_/F_m_ values and pleaching status are highly correlated.

### *Clade C* Symbiodinium *transcriptome*

Although several clade C *Symbiodinium* transcriptomes are publicly available, we assembled a novel transcriptome because our samples were derived from *Siderastrea siderea* and therefore likely divergent from the previously assembled transcriptomes of clade C *Symbiodinium* hosted by the Pacific coral *Acropora hyacinthus* (Barshis et al 2014b) and the Caribbean anemone *Discosoma sanctithomae* (Rosic et al 2015). After adapter trimming and quality filtering, a total of 1,255,626,250 reads were retained (81.5%; 69.6% paired, 11.9% unpaired). The resulting holobiont metatranscriptome contained 333,835 contigs (N50 = 1673), of which 65,838 were unambiguously assigned as *Symbiodinium* specific contigs, with an average length of 1,482 bp and an N50 of 1746. Among symbiont contigs, 45,947 unique isogroups were obtained, of which 22,239 (48.4%) had gene annotations based on sequence homology. Thirty-nine percent of *Symbiodinium* contigs had protein coverage exceeding 0.75 (Supplementary Figure 1) and results from BUSCO suggest that 81.9% of complete and fragmented BUSCOs were present (77.6% complete, 4.3% fragmented), indicating that the transcriptome was fairly comprehensive. All raw reads are archived in the National Center for Biotechnology Information Short Read Archive (SRA) under accession number PRJNA307543, with transcriptome assembly and annotation files available at https://davieslab.wordpress.com/data/ and www.bco-dmo.org/project/635863.

### Symbiodinium *community shifts from clade C to D in response to thermal stress*

Clade-level analysis of sequencing libraries for *Symbiodinium* maintained at the low (25°C) and control (28°C) temperature treatments (for all *p*CO_2_ treatments) revealed that 96.7-100% of reads mapped exclusively to one clade (N=4-436) aligned to clade C *Symbiodinium* (Fig. 2). Analysis of sequencing libraries for *Symbiodinium* maintained at the high temperature treatment (32°C; NS_T32 and FR_T32) consisted of divergent *Symbiodinium* communities hosting 50-75% clade D *Symbiodinium* (Fig. 2). These two libraries were removed from all gene expression analyses due to their divergent clade-level communities, which have been shown to exhibit strong transcriptomic differences (Barshis et al 2014). In addition, these libraries also had the lowest mapped *Symbiodinium* reads (Table 1) due to low sample yield resulting from bleaching under thermal stress.

**Figure 2:**
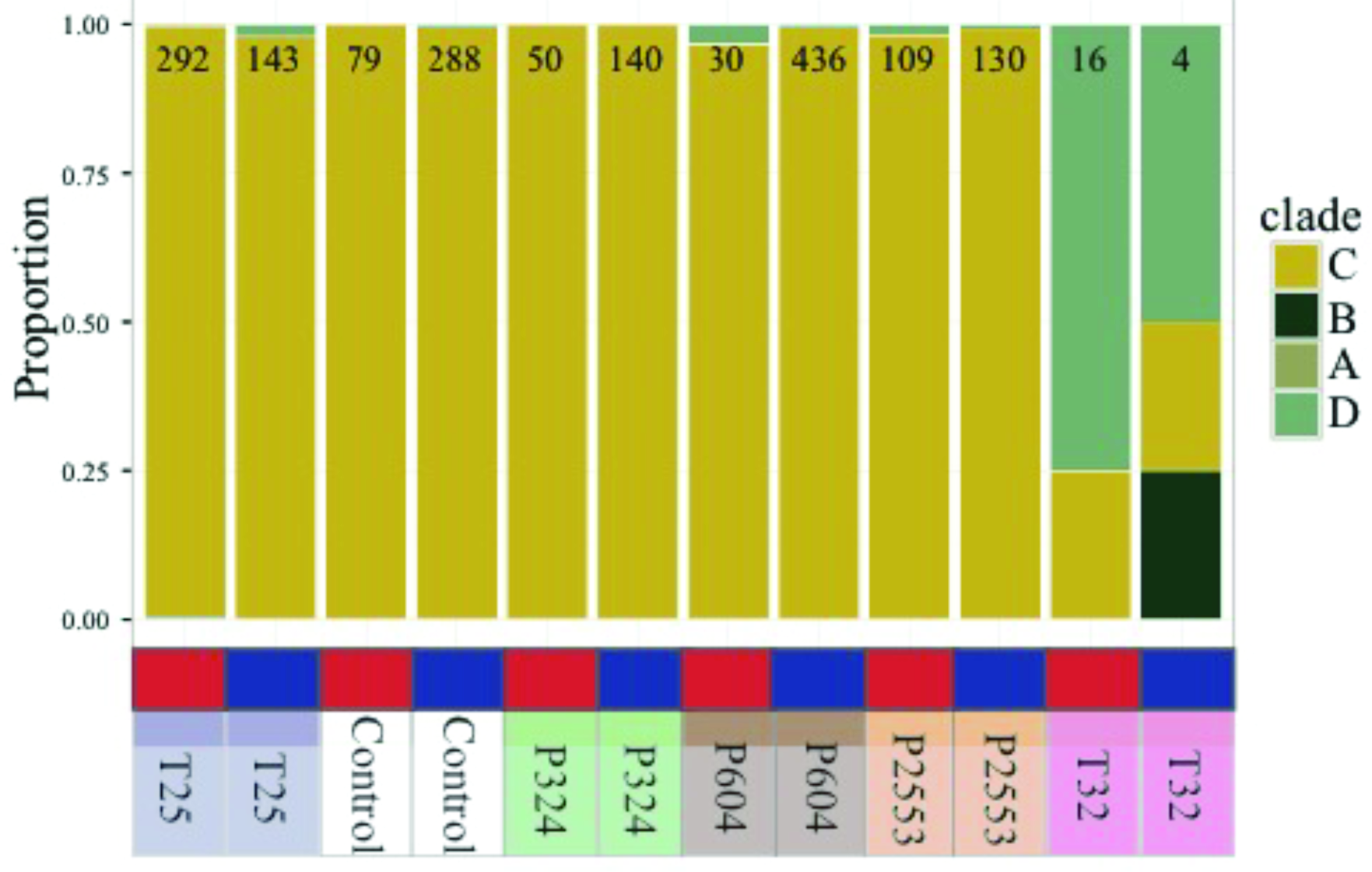
Proportion of RNAseq reads mapping exclusively to one *Symbiodinium* clade lineage (A, B, C, D) for each transcriptome library, with independent columns representing a unique transcriptome library. Number at top of bar indicates total number of reads mapping exclusively to one clade within that library. Red and blue blocks indicate that the library originated from nearshore and forereef corals, respectively. Treatment conditions are noted below the bar (T25= 25.01 ± 0.14 °C, 515 ± 92 μatm; Control= 28.16 ± 0.24 °C, 477 ± 83 μatm; P324= 28.14 ± 0.27 °C, 324 ± 89 μatm; P604= 28.04 ± 0.28 °C, 604 ± 107 μatm; P2553= 27.93 ± 0.19°C, 2553 ± 506 μatm; T32= 32.01 ± 0.17 °C, 472 ± 86 μatm). All libraries were dominated by clade C with the exception of T32 treatments where dominated by clade D.

### Minimal response to acidification stress, with population-specific differences in expression of genes associated with photosynthesis

Without consideration of the high temperature treatment libraries that were excluded due to low mapped reads, transcriptomic analysis of *Symbiodinium* gene expression across the remaining treatments exhibited few differentially expressed genes (DEGs) relative to the control treatment, with 92 (25°C), 40 (324 μatm), 576 (604 μatm) and 23 (2553 μatm)

DEGs observed for the respective treatments, representing a total of 1.69% of the entire transcriptome responding to any treatment (Fig. 3A). In contrast, 24.45% of genes were differentially expressed with respect to reef zone, with 3,792 upregulated in nearshore specimens and 6,441 upregulated in forereef specimens, regardless of treatment condition—demonstrating a strong whole-transcriptomic response to reef zone (Fig. 3B; Fig. 4A, p=0.016). No differences in whole-transcriptome response were detected for treatment (p=0.33; Fig. 3B). Gene ontology enrichment analysis across reef zones revealed many significantly enriched GO terms within cellular components, which were dominated by GO terms associated with photosynthesis in forereef *Symbiodinium* [e.g., *thylakoid part* (GO:0044436), *photosystem* (GO:0009521), *light-harvesting complex* (GO:0030076), *plastid part* (GO:0044435)], suggesting fundamentally different photosynthetic architectures across reef zones (Fig. 4B).

**Figure 3:**
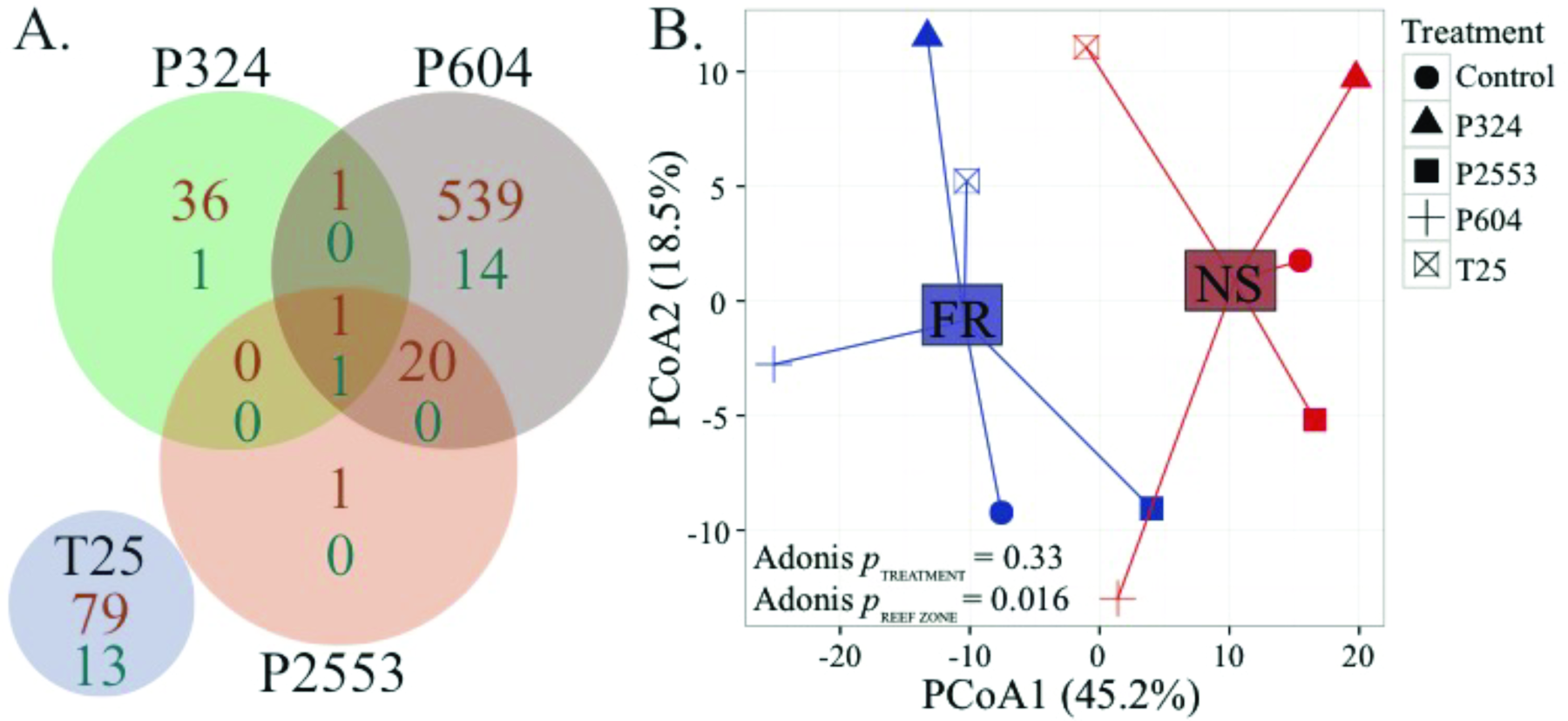
Global RNA-Seq patterns for *in hospite* clade C *Symbiodinium* under different *p*CO_2_ (28 °C) and low temperature (25 °C) treatments. (A) Venn diagram of the number of differentially expressed *Symbiodinium* genes (out of 45,947, FDR=0.05) in experimental conditions relative to control conditions. Orange numbers indicate overrepresented genes and turquoise numbers indicate underrepresented genes, but overall very little transcriptomic response is observed. B: Principal coordinate analysis of all *r-log* transformed isogroups clustered by experimental treatment and reef zone, demonstrating significantly different transcriptome profiles across *Symbiodinium* from different locations, regardless of experimental treatment (NS= nearshore and FR= forereef; T25= 25.01 ± 0.14 °C, 515 ± 92 μatm; Control= 28.16 ± 0.24 °C, 477 ± 83 μatm; P324= 28.14 ± 0.27 °C, 324 ± 89 μatm; P604= 28.04 ± 0.28 °C, 604 ± 107 μatm; P2553= 27.93 ± 0.19°C, 2553 ± 506 μatm).

**Figure 4:**
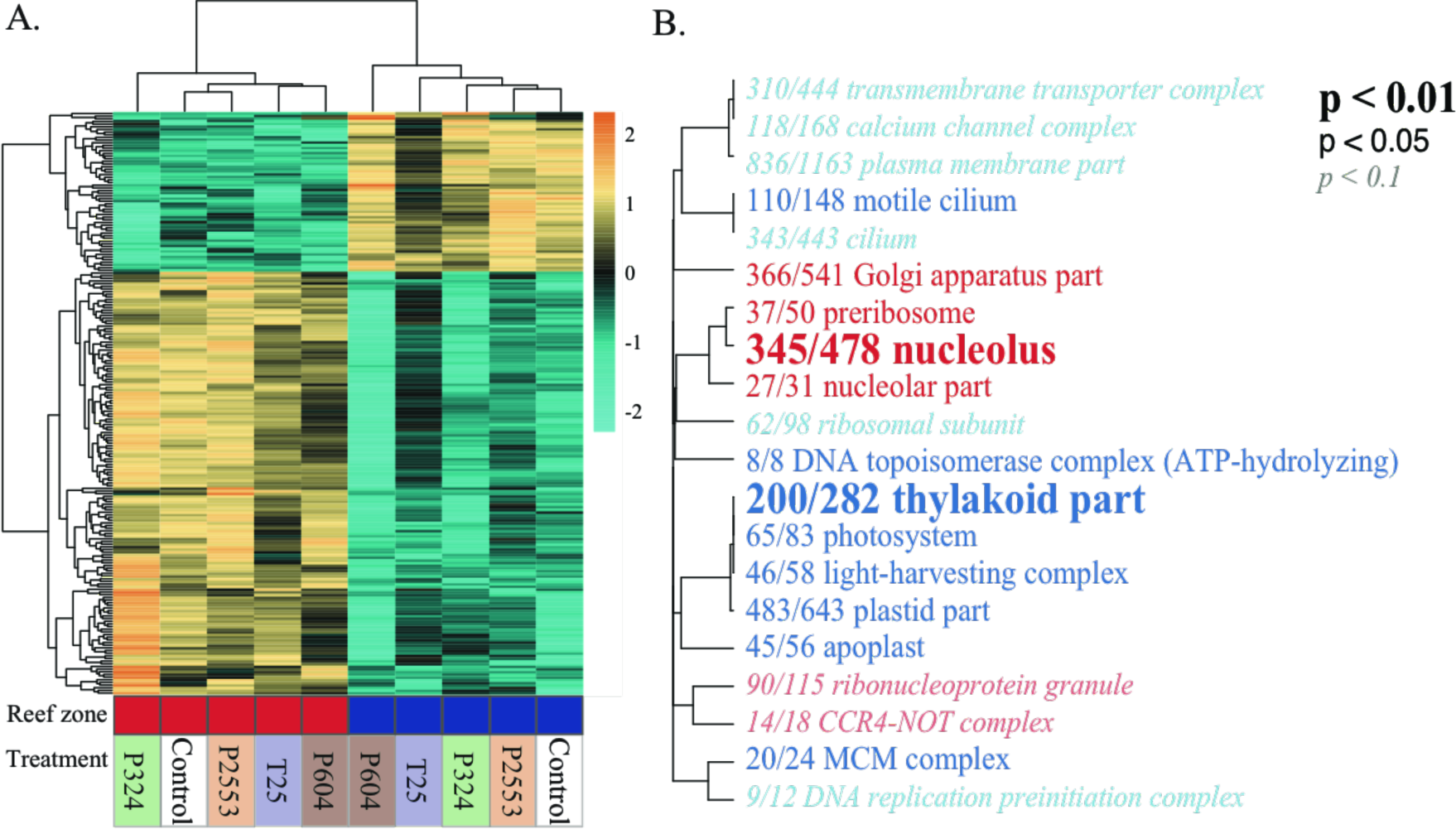
Differentially expressed genes (DEGs) of *Symbiodinium* across reef zones and treatments. (A) Heatmap for the top 300 DEGs for reef zone where each row is a gene and each column is a unique transcriptome library. The color scale is in log2 (fold change relative to the gene’s mean) and genes and samples are clustered hierarchically based on Pearson’s correlation of their expression across samples. Red and blue blocks indicate that libraries originated from nearshore and forereef corals, respectively. Treatment conditions are noted below the bar (T25= 25.01 ± 0.14 °C, 515 ± 92 μatm; Control= 28.16 ± 0.24 °C, 477 ± 83 μatm; P324= 28.14 ± 0.27 °C, 324 ± 89 μatm; P604= 28.04 ± 0.28 °C, 604 ± 107 μatm; P2553= 27.93 ± 0.19°C, 2553 ± 506 μatm). Hierarchical clustering of libraries (columns) demonstrates strong reef zone differences in expression. (B) Gene ontology (GO) enrichment of the ‘Cellular component (CC)’ category derived from the transcriptomic differences across reef zones. Dendrograms depict sharing of genes between categories (categories with no branch length between them are subsets of each other), with the fractions corresponding to proportion of genes with an unadjusted *p* < 0.05 relative to the total number of genes within the category. Text size and boldness indicate the significance (Mann-Whitney U tests) of the term. Blue categories are enriched in forereef *Symbiodinium* while red categories are enriched in the nearshore. Forereef *Symbiodinium* exhibit enrichment of GO categories associated with photosynthesis.

### *Coral host elicits stronger transcriptomic response than* Symbiodinium

When the highly conserved gene (HCG) set from the coral host (*Siderastrea siderea*) and clade C *Symbiodinium* were compared, coral hosts were far more transcriptionally responsive when compared to their *Symbiodinium* partners (Fig. 5). Among these HCGs (N=2,862), corals modified expression by 3.7-15.7% across experimental treatments, while *Symbiodinium* only modified expression by 0.05-2.8%, demonstrating that coral hosts were far more responsive to low temperature and variable acidification stressors when compared to their algal symbionts (Fig. 5). Heatmaps also confirmed that the lack of *Symbiodinium* HCG expression was not driven by strong differences in *Symbiodinium* gene expression across reef zones. Instead genes that were highly differentially expressed in the host showed no differences in expression across reef zones for *Symbiodinium* (Supplementary Fig. S1A-D).

**Figure 5:**
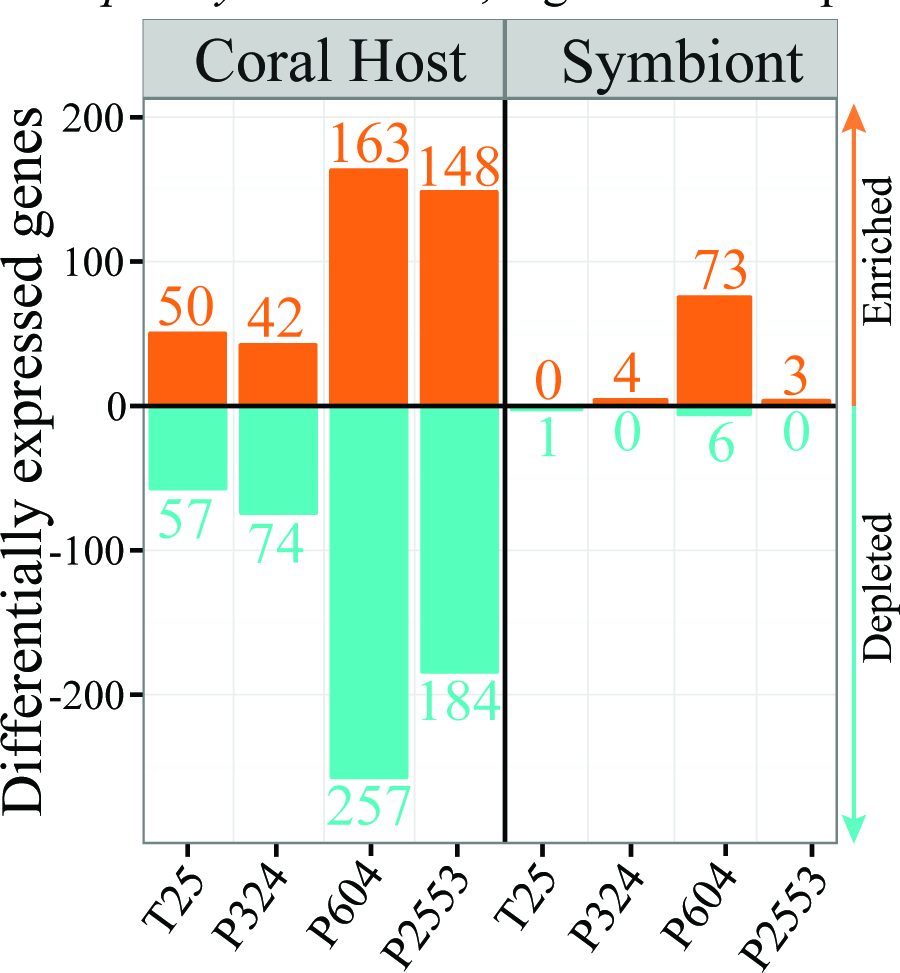
Barplot showing numbers of significantly (*p* < 0.05) differentially expressed genes (DEGs) for the host (left) and its algal symbiont (right) across the four experimental treatments relative to the control for the 2862 highly conserved genes (HCG) across host and symbiont. Orange bars represent enriched genes and turquoise bars represent depleted genes in the treatment (T25= 25.01 ± 0.14 °C, 515 ± 92 μatm; P324= 28.14 ± 0.27 °C, 324 ± 89 μatm; P604= 28.04 ± 0.28 °C, 604 ± 107 μatm; P2553= 27.93 ± 0.19°C, 2553 ± 506 μatm) relative to the control (28.16 ± 0.24 °C, 477 ± 83 μatm). Overall, hosts exhibit a much stronger response across HCGs when compared to *in hospite Symbiodinium*, regardless of experimental treatment.

## DISCUSSION

### *Increased thermotolerance of* Siderastrea siderea *from more thermally variable environments*

Reductions in *Symbiodinium* photosynthetic ability have been strongly correlated with a lack of thermal tolerance and increased susceptibility to bleaching (Howells et al 2012, Takahashi et al 2009, Warner et al 1999). Here we observed strong reductions in photosynthetic efficiency in response to the high temperature treatment (Fig. 1A), which has been consistently observed for *Symbiodinium* exposed to thermal stress (Roth 2014, Warner et al 1996). Strong reductions in calcification rate (Castillo et al 2014) and overall homeostatic disturbance of host gene expression (Davies et al 2016a) was previously observed for these same coral specimens in response to the prescribed thermal stress. These severe reductions in holobiont fitness in response to chronic high temperatures might therefore be driven by the decreases in symbiont photosynthetic function observed here (Fig. 1A).

Notably, we observed that reductions in *S. siderea* symbiont photophysiology at 32°C were dependent on natal reef location, where nearshore *Symbiodinium* were less negatively affected by thermal stress than forereef *Symbiodinium* (Fig. 1A), with photosynthetic efficiency being highly correlated with the coral’s bleaching status (Fig. 1B). Since *Symbiodinium* photophysiology is expected to be a reliable proxy for thermal tolerance (Ragni et al 2010, Wang et al 2012, Warner et al 1996, Warner et al 1999), these fluctuating responses may be due to local adaptation of *Symbiodinium* to more thermally variable nearshore environments (Baumann et al 2016, Castillo and Helmuth 2005). Indeed, Castillo et al. (2012) observed reductions in linear extension rates of forereef *S. siderea* colonies over the past three decades of anthropogenic warming, while nearshore and backreef colonies exhibited constant (nearshore) or increasing (backreef) extension rates over the same interval. Castillo et al (2012) attributed these differential trends in linear extension to thermal preconditioning of backreef and nearshore corals, to the extent that nearshore and backreef environments experience more extreme baseline diurnal and seasonal fluctuations in seawater temperature compared to more thermally stable forereef environments.

The potential for local adaption of *Symbiodinium* is thought to be high given their haploid genomes and short generation times (Correa and Baker 2011, Santos and Coffroth 2003). Howells et al. (2012) observed increased thermotolerance of clade C1 *Symbiodinium* originating from warmer reef locations and Hume et al. (2016) detected strong selection in *S. thermophilium*, a thermotolerant lineage found in extreme temperatures of the Persian Gulf. In addition, recent assisted evolution efforts have shown that *Symbiodinium* can rapidly evolve heat tolerance in culture (Chakravarti et al 2017), providing further evidence of local adaptation in *Symbiodinium*.

### *Chronic high temperatures cause shifts to more thermotolerant* Symbiodinium

*Symbiodinium* genetic diversity has been reported to vary not only among different coral host species and environments but also within a single coral species (Baker 2003, Coffroth and Santos 2005, Thornhill et al 2014, Thornhill et al 2017). This genetic diversity has been shown to drive physiological variation within *Symbiodinium* (Iglesias-Prieto and Trench 1994, Iglesias-Prieto et al 2004, Warner et al 1999) and numerous studies have demonstrated that genetically distinct *Symbiodinium* lineages can exhibit a range of thermotolerances (Robison and Warner 2006, Suggett et al 2008). Consequently, some *Symbiodinium* appear less affected by thermal stress, while others are impacted by even small changes in temperature (Berkelmans and van Oppen 2006, Hume et al 2016, Jones and Berkelmans 2010, Rowan 2004). Here, we observed that long-term thermal stress induced shifts in *Symbiodinium* communities from clade C to D in *S. siderea*, regardless of reef zone origin (Fig. 2). However, this result is at least partially due to a loss of symbiosis with clade C, rather than clade D becoming more successful *in hospite* under increased temperatures, since the corals were still severely bleached (Fig. 1A).

Clade D *Symbiodinium* are generally associated with hosts from marginal reef environments exposed to increased thermal stress (Pettay et al 2011, Pettay and Lajeunesse 2013, Pettay et al 2015) and corals hosting predominantly clade D have been shown to exhibit greater resistance to bleaching (Berkelmans and van Oppen 2006, Jones et al 2008), while those hosting predominantly clade C benefit from higher rates of carbon fixation by the symbionts (Jones and Berkelmans 2010, Stat et al 2008). Although it is unclear from the data at hand if these communities would shift back to clade C dominant communities after alleviation of thermal stress (Sampayo et al 2016), these shifts could represent adaptive potential for corals exposed to future ocean warming (Cunning et al 2015, Silverstein et al 2015).

### Symbiodinium *gene expression and photosynthesis unresponsive to ocean acidification*

Increases in atmospheric *p*CO_2_ have been shown to reduce calcification rates of reef-building corals (e.g., Castillo et al 2014, Chan and Connolly 2013, Hoegh-Guldberg et al 2007, Kleypas et al 1999), but the effects of *p*CO_2_ on the coral’s symbiotic *Symbiodinium* are less clear. Some research suggests that elevated *p*CO_2_ enhances *Symbiodinium* primary production and has proposed that dissolved inorganic carbon is the limiting substrate for photosynthesis, which is more available under high *p*CO2 (Brading et al 2011, Crawley et al 2010). In contrast, others have observed negative effects of increased *p*CO2 on *Symbiodinium* physiology, including reduced productivity, photosynthesis and calcification (Anthony et al 2008, Zhou et al 2016). Here, we observed minimal effects of long-term *p*CO2 stress on *Symbiodinium* photosynthetic efficiencies across a wide range of *p*CO_2_ conditions (Fig. 1A). This lack of response starkly contrasts the strong physiological response and community shift (clade C to D) of *Symbiodinium* under increased temperatures (Figs. 1 & 2). One explanation for this lack of physiological response to *p*CO2 could be that the specific *Symbiodinium* lineage investigated here (clade C) simply does not respond to *p*CO2 stress, which has been observed for other *Symbiodinium* strains (Brading et al 2011).

This lack of physiological response of *Symbiodinium* to acidification stress was also evident at the molecular level, where no significant responses were observed at the whole-transcriptome level (Fig. 3), even when reef zone-specific responses were considered (Fig. S1). This paucity of a transcriptional stress response in *Symbiodinium* is consistent with previous studies, which generally observe little to no transcriptional response to environmental stressors (Barshis et al 2014, Leggat et al 2011, Putnam et al 2013), with the exception of extreme heat stress (Baumgarten et al 2013, Gierz et al 2017, Levin et al 2016)—which we were unable to investigate here due to a combination of severe bleaching (Fig. 1A) and shifts in *Symbiodinium* community (Fig. 2). Instead of responding transcriptionally, it has been proposed that *Symbiodinium* use post-transcriptional regulatory mechanisms, including translational regulation and post-translational modifications, to drive molecular responses. Evidence suggests that very few transcription factors are present in *Symbiodinium* transcriptomes and genomes (Bayer et al 2012, Shoguchi et al 2013). Instead, it has been proposed that these algae utilize small RNAs and microRNAs, (Baumgarten et al 2013, Lin et al 2015), RNA-editing (Liew et al 2017), and trans-splicing of spliced leader sequences (Lin et al 2010, Lin 2011, Zhang et al 2007) to regulate their environmental stress responses. Another possibility is that the timescale of this study (95 days) was not sufficient to trigger physiological and molecular responses in *Symbiodinium*. While the stability of dinoflagellate mRNA is known to be considerably longer than for other organisms (Morey and Van Dolah 2013), it is possible that physiological and transcriptomic responses were minimized by long-term acclimatization of *Symbiodinium* via phenotypic buffering (Reusch 2014). Lastly, and perhaps most likely, it could be that *Symbiodinium in hospite* simply do not respond to changes in *p*CO2 because their position within host-derived tissue-bound space buffers the algae from external changes in pH (Barott et al 2015, Rands et al 1993, Venn et al 2009).

### Functional differences in photosynthesis across divergent environments

*Symbiodinium* exists endosymbiotically across a variety of hosts and habitats, which presents these algae with diverse challenges with respect to photosynthesis and survival. Photosynthetic efficiency of photosystem II (F_v_/F_m_) is widely used as an indicator of photosynthetic performance and stress (Murchie and Lawson 2013). Here, higher F_v_/F_m_ values observed in forereef *Symbiodinium* relative to nearshore (Fig. 1A) might indicate variation in the rate of carbon translocation to hosts (Roberty et al 2014, Warner and Berry-Lowe 2006), since nearshore environments experience increased temperatures and nutrients along with reduced light levels relative to forereef habitats (Baumann et al 2016, Castillo and Lima 2010). These environmental differences may stimulate nearshore corals to rely more heavily on heterotrophy and dissolved/particulate organic matter relative to corals in light-rich forereef habitats (Grottoli et al 2006, Tremblay et al 2014). However, if forereef hosts were receiving increased quantities of photo-assimilates, increased calcification rates would also be expected in forereef corals—a trend that was not previously observed (Castillo et al 2014). Because the physiologies of symbionts from forereef and nearshore corals were divergent even after 95 days in common garden experiments, we propose that the environment has selected for distinct nearshore and forereef *Symbiodinium* ‘ecotypes’ (Chakravarti et al 2017, Howells et al 2012, Iglesias-Prieto et al 2004), although further work is required to test this hypothesis.

Notably, reef-zone-specific photosynthetic efficiencies were also reflected at the transcriptomic level, where fixed differences in gene expression were observed (Fig. 3B, 4A). Divergent stable-state gene expression is perhaps not surprising given that transcriptomic differences across clades (Barshis et al 2014, Rosic et al 2015) and lineages within a clade (Parkinson et al 2016) have been previously observed. Gene ontology enrichment analysis detected strong upregulation of genes related to photosynthesis in forereef symbionts (Fig. 4B), corroborating our observation of higher photosynthetic efficiencies for forereef *Symbiodinium*. These differences in regulation of photosynthetis-related genes could facilitate differences in thermal tolerance across ‘ecotypes’, as was observed here under heat stress (Fig. 1A). Thermally sensitive *Symbiodinium* have been shown to exhibit increased disruption of PSII photochemistry (Robison and Warner 2006, Warner et al 1999), which has been associated with variations in the regulation of genes involved in photosynthesis (McGinley et al 2012). Given that expression differences amongst *Symbiodinium* lineages are consistently enriched for photosynthesis-related genes (Barshis et al 2014, Baumgarten et al 2013, Parkinson et al 2016, Rosic et al 2015), we propose that *Symbiodinium* ‘ecotypes’ have evolved unique transcriptomic signatures for photosynthesis-related genes, which likely reflect functional differences in photosynthetic architectures across environments (Fig. 4B) and represent phenotypic variation upon which selection can act (Chakravarti et al 2017).

### *Coral hosts elicit stronger transcriptomic responses than* Symbiodinium

Understanding how each partner of the coral–*Symbiodinium* symbiosis will respond to environmental stress is required to accurately predict future coral reef susceptibility to global oceanic change (Weis 2008). Although both partners exhibit a wide array of physiological stress responses, *Symbiodinium* are assumed to initiate symbiosis breakdown (Berkelmans and van Oppen 2006, Stat et al 2006, Stat and Gates 2011) due to their production of reactive oxygen species (ROS), which can damage the host (Lesser 1996, Weis 2008). In contrast, the present experiments show that *S. siderea* exhibit stronger transcriptomic responses across highly conserved genes (HCGs) than their *Symbiodinium* symbionts to thermal and CO2-acidification stress (Fig. 5), suggesting that symbiosis breakdown or ‘bleaching’ may be initiated by the host instead of the symbiont. Strong transcriptomic responses of coral hosts are well-documented (Davies et al 2016a, DeSalvo et al 2010b, Meyer et al 2011, Moya et al 2012, Seneca and Palumbi 2015), while *Symbiodinium* transcriptomic responses are generally more subtle (Barshis et al 2014, Baumgarten et al 2013, Leggat et al 2011). For example, Barshis et al. (2014) similarly observed few transcriptional changes across >50,000 genes in response to heat stress across two *Symbiodinium* lineages in symbiosis (D2, C3K), starkly contrasting the broad transcriptomic shifts observed in the symbiont’s host when exposed to identical conditions (Barshis et al 2013). As discussed above, *Symbiodinium* in our study may be unresponsive to the stressors investigated, or the stressors were not applied over a sufficient timescale to elicit a transcriptomic response. Alternatively, the transcriptomic stability of *Symbiodinium* could result from *in hospite* buffering of the symbiosome under *p*CO_2_ stress, which has been observed in response to changes in seawater pH (Barott et al 2015, Rands et al 1993, Venn et al 2009). Although potentially costly to the host, manipulation of *Symbiodinium* responses to stress through active regulation of the symbiont’s environment might be favored over the potential chemical toxicity resulting from the release of reactive molecules by stressed *Symbiodinium* (e.g., Lesser 1996).

### Conclusion

Acclimation and adaptation play critical roles in determining an organism’s ability to tolerate variable environments (Reusch 2014, Schlichting and Pigliucci 1996). Here, we demonstrate that coral-associated *Symbiodinium* exhibit immense capacity for adaptation and acclimation to thermal and acidification stress. *Symbiodinium* from nearshore locations were more resilient to thermal stress than forereef symbionts, suggestive of local adaptation. We also observed differences in gene expression between *Symbiodinium* from nearshore and forereef environments that were coupled with differences in photosynthetic efficiencies, irrespective of treatment condition. These transcriptomic differences suggest that photosynthesis-related gene expression varies by habitat, which likely reflects adaptation to unique environments over potentially long timescales. Host retention of a more thermotolerant clade of *Symbiodinium* (clade D) under thermal stress was also documented, providing further evidence for a mechanism of acclimation to thermal stress. Acclimation to *p*CO_2_ was also observed at both physiological and transcriptomic scales, the mechanisms of which have not yet been identified, although *in hospite* buffering remains a viable hypothesis. Overall, we find evidence for both adaptation and acclimation in the *S. siderea-Symbiodinium* holobiont that may explain the relative resilience of this coral species to global change stressors.

## ACKNOWLEDGEMENTS

We thank Belize Fisheries Department for all associated permits and Garbutt Marine for field assistance. B. Elder, E. Chow, K. Patel, R. Yost, and D. Shroff helped maintain experimental tanks and I. Westfield analyzed carbonate chemistry of the experimental treatments. N. Cohen and K. Delong conducted holobiont tissue preservations and H. Masters performed RNA isolations. Tank experiments were supported by NOAA award NA13OAR4310186 (to JR and KC) and NSF award OCE-1357665 (to JR), sequencing-related activities were supported by AM/KC/JR's start-ups and NSF award 1357665 (to JR), and salary/travel for SD was supported by AM/KC/JR's start-ups, NSF awards OCE-1437371, OCE-1459706 (to JR), and NSF OCE-1459522 (to KC).

---

**Supplemental Figure 1.**
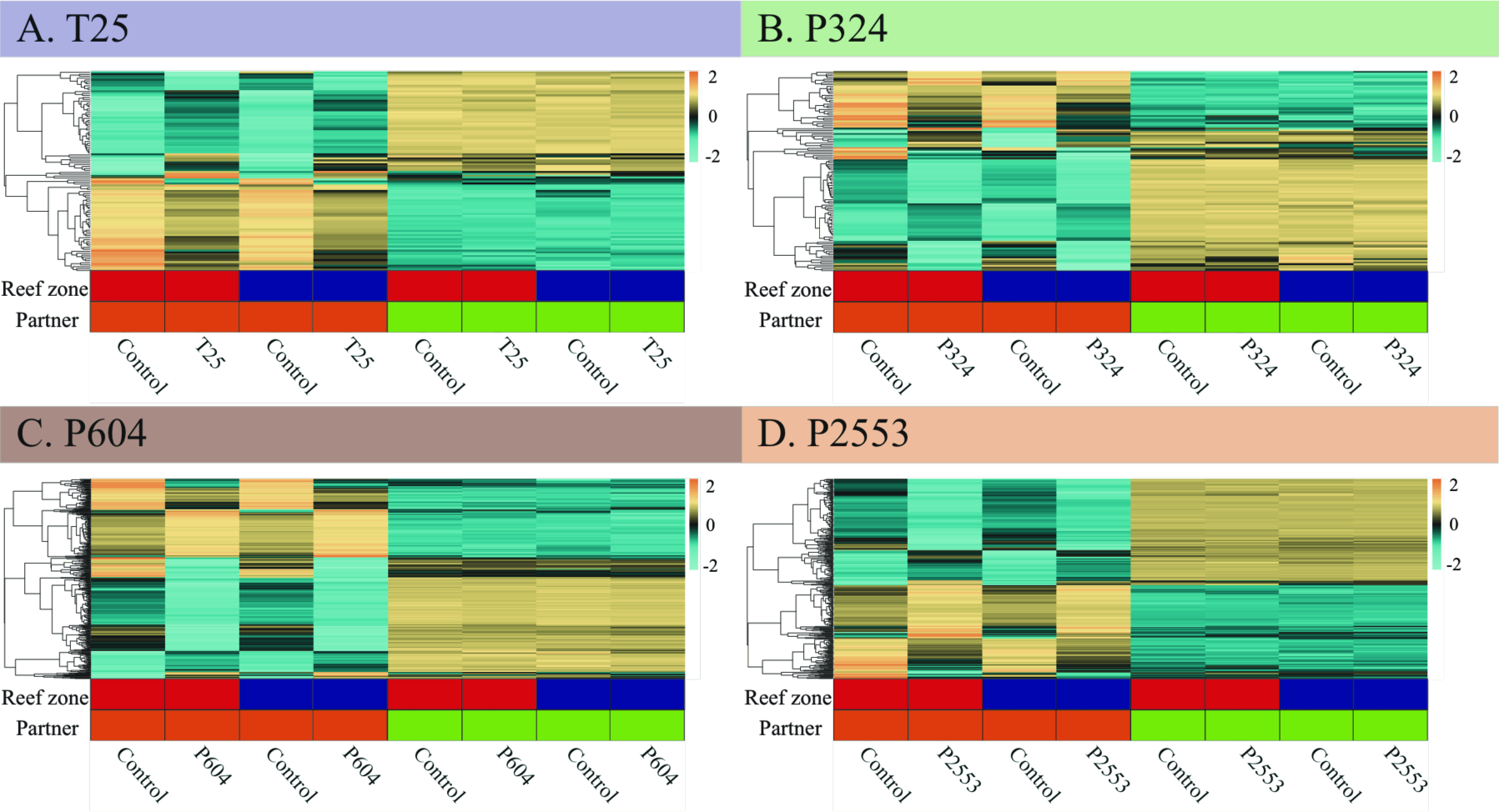
Heatmaps for the expression of highly conserved genes (HCG) found to be differentially expressed in the coral host for each experimental treatment. The left four columns represent expression of these genes in the coral host (orange blocks) and the right four column represent expression of these same genes in *Symbiodinium* (green blocks) across the experimental treatments relative to the control (28.16 ± 0.24 °C, 477 ± 83 μatm): (A) low temperature (T25 = 25.01 ± 0.14 °C, 515 ± 92 μatm), (B) pre-industrial *p*CO_2_ (P324 = 28.14 ± 0.27 °C, 324 ± 89 μatm), (C) next-century *p*CO_2_ (P604 = 28.04 ± 0.28 °C, 604 ± 107 μatm), and (D) extreme-high *p*CO_2_ (P2553 = 27.93 ± 0.19°C, 2553 ± 506 μatm). Red and blue blocks indicate that the library originated from nearshore and forereef reef zones, respectively. Each row is a gene and each column is a unique transcriptome library. The color scale is in log_2_ (fold change relative to the gene’s mean) and genes are clustered hierarchically based on Pearson’s correlation of their expression across samples. Results reveal lack of differential gene expression amongst experimental treatments relative to the control in *Symbiodinium*, compared to strong differential gene expression amongst treatments in the host, and that this result is not due to differential gene expression amongst reef zones in *Symbiodinium*.

## REFERENCES

Anthony KRN, Kline DI, Diaz-Pulido G, Dove S, Hoegh-Guldberg O (2008). Ocean acidification causes bleaching and productivity loss in coral reef builders. P Natl Acad Sci USA 105: 17442-17446.

Baker AC (2001). Ecosystems - Reef corals bleach to survive change. Nature 411: 765-766.

Baker AC (2003). Flexibility and specificity in coral-algal symbiosis: diversity, ecology, and biogeography of Symbiodinium. Annu Rev Ecol Syst 34: 661-689.

Barott KL, Venn AA, Perez SO, Tambutte S, Tresguerres M (2015). Coral host cells acidify symbiotic algal microenvironment to promote photosynthesis. Proc Natl Acad Sci U S A 112: 607-612.

Barshis DJ, Ladner JT, Oliver TA, Seneca FO, Traylor-Knowles N, Palumbi SR (2013). Genomic basis for coral resilience to climate change. Proc Natl Acad Sci U S A 110: 1387-1392.

Barshis DJ, Ladner JT, Oliver TA, Palumbi SR (2014). Lineage-specific transcriptional profiles of Symbiodinium spp. unaltered by heat stress in a coral host. Mol Biol Evol 31: 1343-1352.

Baumann JH, Townsend JE, Courtney TA, Aichelman HE, Davies SW, Lima FP et al (2016). Temperature Regimes Impact Coral Assemblages along Environmental Gradients on Lagoonal Reefs in Belize. Plos One 11.

Baumgarten S, Bayer T, Aranda M, Liew YJ, Carr A, Micklem G et al (2013). Integrating microRNA and mRNA expression profiling in Symbiodinium microadriaticum, a dinoflagellate symbiont of reef-building corals. BMC genomics 14.

Bayer T, Aranda M, Sunagawa S, Yum LK, DeSalvo MK, Lindquist E et al (2012). Symbiodinium Transcriptomes: Genome Insights into the Dinoflagellate Symbionts of Reef-Building Corals. Plos One 7.

Benjamini Y, Hochberg Y (1995). Controlling the false discovery rate: a practical and powerful approach to multiple testing. Journal of the Royal Statistical Society Series B (Methodological): 289-300.

Berkelmans R, van Oppen MJH (2006). The role of zooxanthellae in the thermal tolerance of corals: a 'nugget of hope' for coral reefs in an era of climate change. P Roy Soc B-Biol Sci 273: 2305-2312.

Brading P, Warner ME, Davey P, Smith DJ, Achterberg EP, Suggett DJ (2011). Differential effects of ocean acidification on growth and photosynthesis among phylotypes of Symbiodinium (Dinophyceae). Limnol Oceanogr 56: 927-938.

Castillo KD, Helmuth BST (2005). Influence of thermal history on the response of Montastraea annularis to short-term temperature exposure. Marine Biology 148: 261-270.

Castillo KD, Lima FP (2010). Comparison of in situ and satellite-derived (MODIS-Aqua/Terra) methods for assessing temperatures on coral reefs. Limnol Oceanogr-Meth 8: 107-117.

Castillo KD, Ries JB, Weiss JM, Lima FP (2012). Decline of forereef corals in response to recent warming linked to history of thermal exposure. Nat Clim Change 2: 756-760.

Castillo KD, Ries JB, Bruno JF, Westfield IT (2014). The reef-building coral Siderastrea siderea exhibits parabolic responses to ocean acidification and warming. P Roy Soc B-Biol Sci 281.

Chakravarti LJ, Beltran VH, van Oppen MJH (2017). Rapid thermal adaptation in photosymbionts of reef-building corals. Glob Chang Biol.

Chan NCS, Connolly SR (2013). Sensitivity of coral calcification to ocean acidification: a meta-analysis. Global Change Biol 19: 282-290.

Coffroth MA, Santos SR (2005). Genetic diversity of symbiotic dinoflagellates in the genus Symbiodinium. Protist 156: 19-34.

Consortium TU (2015). UniProt: a hub for protein information. Nucleic Acids Res 43: D204-D212.

Correa AMS, Baker AC (2011). Disaster taxa in microbially mediated metazoans: how endosymbionts and environmental catastrophes influence the adaptive capacity of reef corals. Global Change Biol 17: 68-75.

Crawley A, Kline DI, Dunn S, Anthony K, Dove S (2010). The effect of ocean acidification on symbiont photorespiration and productivity in Acropora formosa. Global Change Biol 16: 851-863.

Cunning R, Silverstein RN, Baker AC (2015). Investigating the causes and consequences of symbiont shuffling in a multi-partner reef coral symbiosis under environmental change. P Roy Soc B-Biol Sci 282.

Davies SW, Marchetti A, Ries JB, Castillo KD (2016a). Thermal and pCO2 Stress Elicit Divergent Transcriptomic Responses in a Resilient Coral. Frontiers in Marine Science 3.

Davies SW, Wham D, Kanke MR, Matz MV (2016b). Ecological factors rather than barriers to dispersal shape genetic structure of algal symbionts in horizontally-transmitting corals. bioRxiv.

DeSalvo MK, Sunagawa S, Fisher PL, Voolstra CR, Iglesias-Prieto R, Medina M (2010a). Coral host transcriptomic states are correlated with Symbiodinium genotypes. Mol Ecol 19: 1174-1186.

DeSalvo MK, Sunagawa S, Voolstra CR, Medina M (2010b). Transcriptomic responses to heat stress and bleaching in the elkhorn coral Acropora palmata. Mar Ecol Prog Ser 402: 97-113.

Dixon GB, Davies SW, Aglyamova GA, Meyer E, Bay LK, Matz MV (2015). Genomic determinants of coral heat tolerance across latitudes. Science 348: 1460-1462.

Franklin EC, Stat M, Pochon X, Putnam HM, Gates RD (2012). GeoSymbio: a hybrid, cloud-based web application of global geospatial bioinformatics and ecoinformatics for Symbiodinium-host symbioses. Molecular ecology resources 12: 369-373.

Gardner SG, Raina JB, Ralph PJ, Petrou K (2017). Reactive oxygen species (ROS) and dimethylated sulphur compounds in coral explants under acute thermal stress. The Journal of experimental biology.

Gierz SL, Foret S, Leggat W (2017). Transcriptomic Analysis of Thermally Stressed Symbiodinium Reveals Differential Expression of Stress and Metabolism Genes. Front Plant Sci 8.

Grabherr MG, Haas BJ, Yassour M, Levin JZ, Thompson DA, Amit I et al (2011). Full-length transcriptome assembly from RNA-Seq data without a reference genome. Nat Biotechnol 29: 644-U130.

Grottoli AG, Rodrigues LJ, Palardy JE (2006). Heterotrophic plasticity and resilience in bleached corals. Nature 440: 1186-1189.

Hoegh-Guldberg O (1999). Climate change, coral bleaching and the future of the world's coral reefs. Marine and Freshwater Research 50: 839-866.

Hoegh-Guldberg O, Mumby PJ, Hooten AJ, Steneck RS, Greenfield P, Gomez E et al (2007). Coral reefs under rapid climate change and ocean acidification. Science 318: 1737-1742.

Hoegh-Guldberg O, Bruno JF (2010). The impact of climate change on the world's marine ecosystems. Science 328: 1523-1528.

Howells EJ, Beltran VH, Larsen NW, Bay LK, Willis B, van Oppen MJ (2012). Coral thermal tolerance shaped by local adaptation of photosymbionts. Nat Clim Change 2: 116-120.

Hughes TP, Kerry JT, Alvarez-Noriega M, Alvarez-Romero JG, Anderson KD, Baird AH et al (2017). Global warming and recurrent mass bleaching of corals. Nature 543: 373-377.

Hume BCC, Voolstra CR, Arif C, D'Angelo C, Burt JA, Eyal G et al (2016). Ancestral genetic diversity associated with the rapid spread of stress-tolerant coral symbionts in response to Holocene climate change. P Natl Acad Sci USA 113: 4416-4421.

Iglesias-Prieto R, Matta JL, Robins WA, Trench RK (1992). Photosynthetic Response to Elevated-Temperature in the Symbiotic Dinoflagellate Symbiodinium-Microadriaticum in Culture. P Natl Acad Sci USA 89: 10302-10305.

Iglesias-Prieto R, Trench RK (1994). Acclimation and Adaptation to Irradiance in Symbiotic Dinoflagellates. 1. Responses of the Photosynthetic Unit to Changes in Photon Flux-Density. Mar Ecol Prog Ser 113: 163-175.

Iglesias-Prieto R, Beltran VH, LaJeunesse TC, Reyes-Bonilla H, Thome PE (2004). Different algal symbionts explain the vertical distribution of dominant reef corals in the eastern Pacific. Proceedings Biological sciences / The Royal Society 271: 1757-1763.

Jones A, Berkelmans R (2010). Potential costs of acclimatization to a warmer climate: growth of a reef coral with heat tolerant vs. sensitive symbiont types. Plos One 5: e10437.

Jones AM, Berkelmans R, van Oppen MJH, Mieog JC, Sinclair W (2008). A community change in the algal endosymbionts of a scleractinian coral following a natural bleaching event: field evidence of acclimatization. P Roy Soc B-Biol Sci 275: 1359-1365.

Kemp DW, Hernandez-Pech X, Iglesias-Prieto R, Fitt WK, Schmidt GW (2014). Community dynamics and physiology of Symbiodinium spp. before, during, and after a coral bleaching event. Limnol Oceanogr 59: 788-797.

Kleypas JA, Buddemeier RW, Archer D, Gattuso JP, Langdon C, Opdyke BN (1999). Geochemical consequences of increased atmospheric carbon dioxide on coral reefs. Science 284: 118-120.

Kolde R (2012). pheatmap: Pretty Heatmaps. R package version 0.6.1. [http://CRANR-projectorg/package=pheatmap%5D

Langmead B, Salzberg SL (2012). Fast gapped-read alignment with Bowtie 2. Nat Methods 9: 357-U354.

Leggat W, Seneca F, Wasmund K, Ukani L, Yellowlees D, Ainsworth TD (2011). Differential Responses of the Coral Host and Their Algal Symbiont to Thermal Stress. Plos One 6.

Lesser MP (1996). Acclimation of phytoplankton to UV-B radiation: Oxidative stress and photoinhibition of photosynthesis are not prevented by UV-absorbing compounds in the dinoflagellate Prorocentrum micans. Mar Ecol Prog Ser 132: 287-297.

Lesser MP, Stat M, Gates RD (2013). The endosymbiotic dinoflagellates (Symbiodinium sp.) of corals are parasites and mutualists. Coral Reefs: 603-611.

Levin RA, Beltran VH, Hill R, Kjelleberg S, McDougald D, Steinberg PD et al (2016). Sex, Scavengers, and Chaperones: Transcriptome Secrets of Divergent Symbiodinium Thermal Tolerances (vol 33, pg 2201, 2016). Molecular Biology and Evolution 33: 3032-3032.

Liew YJ, Li Y, Baumgarten S, Voolstra CR, Aranda M (2017). Condition-specific RNA editing in the coral symbiont Symbiodinium microadriaticum. Plos Genet 13.

Lin SJ, Zhang HA, Zhuang YY, Tran B, Gill J (2010). Spliced leader-based metatranscriptomic analyses lead to recognition of hidden genomic features in dinoflagellates. P Natl Acad Sci USA 107: 20033-20038.

Lin SJ (2011). Genomic understanding of dinoflagellates. Res Microbiol 162: 551-569.

Lin SJ, Cheng SF, Song B, Zhong X, Lin X, Li WJ et al (2015). The Symbiodinium kawagutii genome illuminates dinoflagellate gene expression and coral symbiosis. Science 350: 691-694.

Love MI, Huber W, Anders S (2014). Moderated estimation of fold change and dispersion for RNA-seq data with DESeq2. Genome Biol 15: 550.

Martin JA, Wang Z (2011). Next-generation transcriptome assembly. Nature reviews Genetics 12: 671-682.

McGinley MP, Aschaffenburg MD, Pettay DT, Smith RT, LaJeunesse TC, Warner ME (2012). Transcriptional Response of Two Core Photosystem Genes in Symbiodinium spp. Exposed to Thermal Stress. Plos One 7.

Meyer E, Aglyamova GV, Matz MV (2011). Profiling gene expression responses of coral larvae (Acropora millepora) to elevated temperature and settlement inducers using a novel RNA-Seq procedure. Mol Ecol 20: 3599-3616.

Morey JS, Van Dolah FM (2013). Global analysis of mRNA half-lives and de novo transcription in a dinoflagellate, Karenia brevis. Plos One 8: e66347.

Moya A, Huisman L, Ball EE, Hayward DC, Grasso LC, Chua CM et al (2012). Whole transcriptome analysis of the coral Acropora millepora reveals complex responses to CO(2)-driven acidification during the initiation of calcification. Mol Ecol 21: 2440-2454.

Murchie EH, Lawson T (2013). Chlorophyll fluorescence analysis: a guide to good practice and understanding some new applications. J Exp Bot 64: 3983-3998.

Muscatine L, Cernichiari E (1969). Assimilation of photosynthetic products of zooxanthellae by a reef coral. The Biological Bulletin 137: 506-523.

Muscatine L (1990). The role of symbiotic algae in carbon and energy flux in reef corals. Ecosystems of the World 25: 75-87.

Oksanen JF, Blanchet G, Kindt R, Legendre P, Minchin PR, O'Hara RB et al (2013). vegan: Community Ecology Package. R package version 2.0-7.

Pandolfi JM, Connolly SR, Marshall DJ, Cohen AL (2011). Projecting Coral Reef Futures Under Global Warming and Ocean Acidification. Science 333: 418-422.

Parkinson JE, Baumgarten S, Michell CT, Baums IB, LaJeunesse TC, Voolstra CR (2016). Gene Expression Variation Resolves Species and Individual Strains among Coral-Associated Dinoflagellates within the Genus Symbiodinium. Genome Biology and Evolution 8: 665-680.

Pettay DT, Wham DC, Pinzon JH, LaJeunesse TC (2011). Genotypic diversity and spatial-temporal distribution of Symbiodinium clones in an abundant reef coral. Mol Ecol 20: 5197-5212.

Pettay DT, Lajeunesse TC (2013). Long-range dispersal and high-latitude environments influence the population structure of a "stress-tolerant" dinoflagellate endosymbiont. Plos One 8: e79208.

Pettay DT, Wham DC, Smith RT, Iglesias-Prieto R, LaJeunesse TC (2015). Microbial invasion of the Caribbean by an Indo-Pacific coral zooxanthella. P Natl Acad Sci USA 112: 7513-7518.

Pochon X, Montoya-Burgos JI, Stadelmann B, Pawlowski J (2006). Molecular phylogeny, evolutionary rates, and divergence timing of the symbiotic dinoflagellate genus Symbiodinium. Molecular phylogenetics and evolution 38: 20-30.

Pochon X, Gates RD (2010). A new Symbiodinium clade (Dinophyceae) from soritid foraminifera in Hawai'i. Molecular phylogenetics and evolution 56: 492-497.

Putnam HM, Mayfield AB, Fan TY, Chen CS, Gates RD (2013). The physiological and molecular responses of larvae from the reef-building coral Pocillopora damicornis exposed to near-future increases in temperature and pCO(2). Marine Biology 160: 2157-2173.

R Development Core Team (2015). R: A language and environment for statistical computing. R Foundation for Statistical Computing: Vienna, Austria.

Ragni M, Airs RL, Hennige SJ, Suggett DJ, Warner ME, Geider RJ (2010). PSII photoinhibition and photorepair in Symbiodinium (Pyrrhophyta) differs between thermally tolerant and sensitive phylotypes. Mar Ecol Prog Ser 406: 57-70.

Rands ML, Loughman BC, Douglas AE (1993). The Symbiotic Interface in an Alga Invertebrate Symbiosis. P Roy Soc B-Biol Sci 253: 161-165.

Reusch TB (2014). Climate change in the oceans: evolutionary versus phenotypically plastic responses of marine animals and plants. Evol Appl 7: 104-122.

Roberty S, Bailleul B, Berne N, Franck F, Cardol P (2014). PSI Mehler reaction is the main alternative photosynthetic electron pathway in Symbiodinium sp., symbiotic dinoflagellates of cnidarians. New Phytol 204: 81-91.

Robison JD, Warner ME (2006). Differential impacts of photoacclimation and thermal stress on the photobiology of four different phylotypes of Symbiodinium (Pyrrhophyta). J Phycol 42: 568-579.

Rosic N, Ling EY, Chan CK, Lee HC, Kaniewska P, Edwards D et al (2015). Unfolding the secrets of coral-algal symbiosis. Isme J 9: 844-856.

Roth MS (2014). The engine of the reef: photobiology of the coral-algal symbiosis. Front Microbiol 5.

Rowan R (2004). Coral bleaching - Thermal adaptation in reef coral symbionts. Nature 430: 742-742.

Sampayo EM, Ridgway T, Franceschinis L, Roff G, Hoegh-Guldberg O, Dove S (2016). Coral symbioses under prolonged environmental change: living near tolerance range limits. Scientific reports 6.

Santos SR, Coffroth MA (2003). Molecular Genetic Evidence that Dinoflagellates Belonging to the Genus Symbiodinium Freudenthal Are Haploid. Biol Bull 204: 10-20.

Schlichting C, Pigliucci M (1996). Phenotypic Evolution: A Reaction Norm Perspective․ ․ Sinauer Associates: Sunderland, MA.

Seneca FO, Palumbi SR (2015). The role of transcriptome resilience in resistance of corals to bleaching. Mol Ecol 24: 1467-1484.

Shoguchi E, Shinzato C, Kawashima T, Gyoja F, Mungpakdee S, Koyanagi R et al (2013). Draft assembly of the Symbiodinium minutum nuclear genome reveals dinoflagellate gene structure. Curr Biol 23: 1399-1408.

Silverstein RN, Cunning R, Baker AC (2015). Change in algal symbiont communities after bleaching, not prior heat exposure, increases heat tolerance of reef corals. Global Change Biol 21: 236-249.

Simao FA, Waterhouse RM, Ioannidis P, Kriventseva EV, Zdobnov EM (2015). BUSCO: assessing genome assembly and annotation completeness with single-copy orthologs. Bioinformatics 31: 3210-3212.

Stat M, Carter D, Hoegh-Guldberg O (2006). The evolutionary history of Symbiodinium and scleractinian hosts - Symbiosis, diversity, and the effect of climate change. Perspect Plant Ecol 8: 23-43.

Stat M, Morris E, Gates RD (2008). Functional diversity in coral-dinoflagellate symbiosis. Proc Natl Acad Sci U S A 105: 9256-9261.

Stat M, Gates RD (2011). Clade D Symbiodinum in scleractinian corals: A "nugget" of hope, a selfish opportunist, an ominous sign, or all of the above. Marine Biology 730315: 1-9.

Suggett DJ, Warner ME, Smith DJ, Davey P, Hennige S, Baker NR (2008). Photosynthesis and production of hydrogen peroxide by Symbiodinium (Pyrrhophyta) phylotypes with different thermal tolerances. J Phycol 44: 948-956.

Takahashi S, Whitney SM, Badger MR (2009). Different thermal sensitivity of the repair of photodamaged photosynthetic machinery in cultured Symbiodinium species. P Natl Acad Sci USA 106: 3237-3242.

Thornhill DJ, LaJeunesse TC, Kemp DW, Fitt WK, Schmidt GW (2006). Multi-year, seasonal genotypic surveys of coral-algal symbioses reveal prevalent stability or post-bleaching reversion. Marine Biology 148: 711-722.

Thornhill DJ, Lewis AM, Wham DC, LaJeunesse TC (2014). Host-specialist lineages dominate the adaptive radiation of reef coral endosymbionts. Evolution 68: 352-367.

Thornhill DJ, Howells EJ, Wham DC, Steury TD, Santos SR (2017). Population genetics of reef coral endosymbionts (Symbiodinium, Dinophyceae). Mol Ecol.

Tremblay P, Fine M, Maguer JF, Grover R, Ferrier-Pages C (2013). Photosynthate translocation increases in response to low seawater pH in a coral-dinoflagellate symbiosis. Biogeosciences 10: 3997-4007.

Tremblay P, Grover R, Maguer JF, Hoogenboom M, Ferrier-Pages C (2014). Carbon translocation from symbiont to host depends on irradiance and food availability in the tropical coral Stylophora pistillata. Coral Reefs 33: 1-13.

Trench RK, Blank RJ (1987). Symbiodinium microadriaticum Fredenthal, S. Goreau sp. nov., S. Kawagut2 sp. nov., and S. pilosum sp. nov.: Gymnodinoid dinoflagellate symbionts of marine invertebrates. J Phycol 23: 469-481.

Venn AA, Tambutte E, Lotto S, Zoccola D, Allemand D, Tambutte S (2009). Imaging intracellular pH in a reef coral and symbiotic anemone. Proc Natl Acad Sci U S A 106: 16574-16579.

Voolstra CR, Sunagawa S, Matz MV, Bayer T, Aranda M, Buschiazzo E et al (2011). Rapid evolution of coral proteins responsible for interaction with the environment. Plos One 6: e20392.

Wang JT, Meng PJ, Chen YY, Chen CA (2012). Determination of the Thermal Tolerance of Symbiodinium Using the Activation Energy for Inhibiting Photosystem II Activity. Zool Stud 51: 137-142.

Warner ME, Fitt WK, Schmidt GW (1996). The effects of elevated temperature on the photosynthetic efficiency of zooxanthellae in hospite from four different species of reef coral: A novel approach. Plant Cell and Environment 19: 291-299.

Warner ME, Fitt WK, Schmidt GW (1999). Damage to photosystem II in symbiotic dinoflagellates: A determinant of coral bleaching. P Natl Acad Sci USA 96: 8007-8012.

Warner ME, Berry-Lowe S (2006). Differential xanthophyll cycling and photochemical activity in symbiotic dinoflagellates in multiple locations of three species of Caribbean coral. J Exp Mar Biol Ecol 339: 86-95.

Weis VM (2008). Cellular mechanisms of Cnidarian bleaching: stress causes the collapse of symbiosis. Journal of Experimental Biology 211: 3059-3066.

Winters G, Holzman R, Blekhman A, Beer S, Loya Y (2009). Photographic assessment of coral chlorophyll contents: Implications for ecophysiological studies and coral monitoring. J Exp Mar Biol Ecol 380: 25-35.

Yellowlees D, Rees TA, Leggat W (2008). Metabolic interactions between algal symbionts and invertebrate hosts. Plant, cell & environment 31: 679-694.

Zhang H, Hou YB, Miranda L, Campbell DA, Sturm NR, Gaasterland T et al (2007). Spliced leader RNA trans-splicing in dinoflagellates. P Natl Acad Sci USA 104: 46184623.

Zhou GW, Yuan T, Cai L, Zhang WP, Tian RM, Tong HY et al (2016). Changes in microbial communities, photosynthesis and calcification of the coral Acropora gemmifera in response to ocean acidification. Scientific reports 6.

